# Modelling Glomerular Filtration using Multispectral Optoacoustic Tomography and a Novel Near-infrared Dye

**DOI:** 10.1101/2025.05.31.657147

**Authors:** James Littlewood, Rachel Harwood, Srishti Vajpayee, Rossana Perciaccante, Norbert Gretz, Katherine Trivino-Cepeda, Jack Sharkey, Thomas Sardella, Neal Burton, Arthur Taylor, Bettina Wilm, Rachel Bearon, Patricia Murray

**Affiliations:** Institute of Life Course and Medical Sciences, University of Liverpool, Liverpool, L69 3BX, United Kingdom; iThera Medical, Zielstattstraße 13, 81379 Munich, Germany; Cyanagen Srl, Via degli Stradelli Guelfi 40/C, 40138 Bologna, Italy; Heidelberg University, Grabengasse 1, Heidelberg, DE 69117, Germany; Revvity Inc, Chalfont Road, Seer Green, Beaconsfield, HP9 2FX, United Kingdom; Dept of Mathematical Sciences, University of Liverpool, Liverpool, L69 3BX, United Kingdom

**Keywords:** Multispectral optoacoustic tomography, near-infrared dyes, kidney function, glomerular filtration rate, ischaemia reperfusion injury, mouse model

## Abstract

Kidney disease carries significant morbidity, mortality, and financial cost. Preclinical work exploring more accurate methods for assessing renal function is key for monitoring disease progression and the efficacy of future therapies. Measuring kidney function by dynamic contrast enhanced imaging allows us to measure left and right kidney function separately. This provides an internal control where a unilateral injury model is used. Multispectral optoacoustic tomography (MSOT) can measure the renal clearance of administered near infrared absorbers and, via the application of mathematical models, allows for the calculation of glomerular filtration rate. ABZWCY-HPβCD is a cyanine dye bound to 2-hydroxylpropyl-β-cyclodextrin and has shown promise as a glomerular filtration marker. Here we combine MSOT, ABZWCY-HPβCD, and a modified Patlak-Rutland model to measure the glomerular filtration rate of healthy mice and following ischaemic reperfusion injury.

## 1 Introduction

Kidney disease carries significant morbidity, mortality, and financial cost and its prevalence is rising [1–8]. Rodent kidney models, such as the ischaemia-reperfusion injury (IRI) model, are invaluable for investigating kidney disease and have previously facilitated a deep understanding of the molecular mechanisms of acute kidney injury (AKI) and chronic kidney disease (CKD) [9–13]. IRI involves the restriction and then restoration of blood flow to a tissue which can be achieved in animals by applying a surgical clamp to an artery. Both the ischaemic and reperfusion phases contribute to injury [14]. IRI has significant clinical relevance as it occurs during solid organ transplantation, angioplasty, cardiac bypass, and tissue hypoperfusion (thrombosis, haemorrhage, allergy, gastrointestinal fluid losses, heart failure, and sepsis amongst others) [14–16].

An accurate measurement of kidney function is important in preclinical experiments, with non-invasive techniques that can be used longitudinally being particularly advantageous [17,18]. The glomerular filtration rate (GFR) is the current gold standard measurement of renal function in the clinic. GFR correlates with the amount of functioning renal tissue and the ability of the kidneys to maintain fluid and electrolyte homeostasis, produce hormones, and clear nitrogenous waste [19,20]. Serum creatinine is often used in humans to estimate GFR, but it is not a very accurate indicator of GFR in mice due to up to 50% of creatinine being secreted by the tubules of the rodent nephron [21]. Measuring any blood marker repeatedly in mice is undesirable due to their small total blood volume, meaning sequential samples risk hypovolaemia which alters GFR and negatively impacts animal welfare. The GFR marker inulin bound to a fluorescent label has been injected into mice and measured in the urine to quantify GFR [22]. This requires placing the mouse in a metabolic cage that adds further potential stress for the mice as well as evidence of altered renal function [23]. Improving on this, transcutaneous optical devices have been used to measure the GFR markers inulin, sinistrin, and cyclodextrin via a fluorescent label and model their disappearance from the skin to quantify GFR [24–28].

Measuring kidney function by dynamic contrast enhanced (DCE) imaging offers the distinct benefit of separately measuring left and right kidney function which can differ even in healthy animals [29,30]. Additionally, measuring both kidneys where a unilateral injury has been applied offers the advantage of an internal control. Several imaging modalities have been used to measure glomerular filtration rate (GFR) with DCE, including magnetic resonance imaging (MRI) [31–34], computed tomography (CT) [35–37], and the nuclear medicine techniques of scintigraphy and single-photon emission computed tomography (SPECT) [38–41]. DCE MRI to measure GFR has been limited by balancing the trade-off between spatial and temporal resolution, uncertainty about accuracy [31,42–45], and the risk of nephrogenic systemic fibrosis with gadolinium contrast agents [46]. Nuclear medicine techniques require operators to work with harmful radioactive materials and expose the subject to ionising radiation. Candidate CT GFR contrast agents e.g. iothalamate have their controversies including nephrotoxicity at high osmolarity [47] and evidence of tubular secretion [48] or reabsorption [49].

Multispectral optoacoustic tomography (MSOT) can measure renal function noninvasively and longitudinally without radiation and at a simultaneously high spatial and temporal resolution. Measuring GFR by MSOT requires an appropriate contrast agent. Jiang et al have measured SK-GFR with MSOT via the renal clearance of a gold nanocluster, Au_25_(SG)_18_, in combination with a Patlak-Rutland model [50]. This was examined in both healthy mice and following a surgical injury model of unilateral ureteral obstruction. However, alternative contrast agents for measuring GFR in MSOT are worth exploring as gold nanoparticles have a higher risk of toxicity, show poor biodegradability, and are relatively expensive [51–53]. Toxicity is a particular concern for in vivo studies due to the gold nanoparticles having toxic effects on organs such as the liver and kidneys [54].

Small-molecule organic near-infrared (NIR) dyes show potential benefits in MSOT. They are associated with low toxicity, high biocompatibility, and rapid clearance [51,55]. They can be designed to have a variety of ionic charges and different degrees of protein binding that results in their potential clearance by different excretory routes in vivo including glomerular filtration and tubular secretion.

Several studies have used preclinical MSOT to assess renal clearance by measuring the kinetics of the NIR dye IRDye 800CW [17,18,56–62]. The biodistribution of IRDye 800CW in rodents is primarily in the kidneys and urine, but it is also present in the liver and other organs [63]. Studies using IRDye 800CW have shown correlation with kidney injury in rodent models of IRI [56], diabetic kidney disease [59], septic acute kidney injury [57], and primary focal segmental glomerulosclerosis [17,18]. However, the specificity of IRDye 800CW for assessing renal function is in question due to its high levels of plasma protein binding (41%) [17] and evidence of the dye in the extrahepatic bile ducts of live rodents after intravenous administration [64]. As such, IRDye 800CW cannot be used to accurately calculate GFR.

ABZWCY-HPβCD is a zwitterionic heptamethine cyanine NIR dye bound to 2-hydroxylpropyl-β-cyclodextrin with a peak optical absorbance of 705 nm and a molecular weight of 2,466 Da [28,65]. ABZWCY-HPβCD has shown great potential as a marker of GFR in rodents [28], but it has not previously been examined using MSOT. Huang et al have shown that ABZWCY-HPβCD has low plasma protein binding (3.7%), has high levels of urinary recovery (97% after 24 hours), does not undergo tubular secretion, and is not cytotoxic on incubation with human proximal tubular cells [28]. The urinary recovery of ABZWCY-HPβCD compares favourably with Au_25_(SG)_18_ (70% after 24 hours) [50].

The aim of this study was to assess the suitability of ABZWCY-HPβCD as a dynamic contrast agent using MSOT and apply an appropriate mathematical model to measure single kidney glomerular filtration rate (SK-GFR).

## 2 Materials and methods

### 2.1 Dye Solutions

Unless otherwise specified, compounds were dissolved in sterile 1X Dulbecco′s Phosphate Buffered Saline (PBS) (Sigma, UK) and thoroughly mixed by vortexing. Solutions were passed through 0.2 μm sterile filters (Sartorius, Germany) and stored in microfuge tubes at -20°C and wrapped in tin foil. ABZWCY-HPβCD was provided by Cyanagen, Italy. This product is now marketed as STAR FLUOR 770 – GFR. IRDye 800CW Carboxylate was purchased from LI-COR, USA. Fluorescein isothiocyanate-sinistrin (FITC-sinistrin) was purchased from MediBeacon, Germany.

### 2.2 Photospectra Measurements

Photospectra of dyes was measured by the FLUOstar Omega microplate reader (BMG LABTECH, France) in flat bottomed polystyrene “costar” 96-well microplates (Corning, USA). Automatic path correction was applied and a volume of 100 μL was used. A separate set of control wells containing the solvents was measured and the absorbance was subtracted from the absorbance of the dye wells. Wavelengths from 680 to 900 nm were measured in steps of 1 nm.

### 2.3 Agar Phantom Creation

Tissue mimicking, scattering (μ′s = 5 cm^-1^), non-absorbing phantoms of 2 cm diameter were made by a previously published method [65–67]. The light scattering property, which is similar to living tissue, is provided by intralipid (a lipid emulsion from soybeans). A mould was made by cutting off the front of a 20 mL, 2 cm diameter syringe. 1.03 mL of 20% intralipid (Sigma-Aldrich, USA) was warmed in a water bath at 60°C. Separately, 0.75 g of agar (Sigma, UK) was added to 50 mL of distilled water and heated until boiling in a microwave. The warmed intralipid was added to the hot agar and stirred, then poured into the mould. Two straws of 4 mm bore (Stephensons, UK) were inserted before the solution could set and positioned using tape. These created cavities for insertions of test absorbers later. The phantoms were allowed to set at room temperature for one hour.

### 2.4 Animals

All animal work was carried out by Home Office Personal Licence holders under the Home Office Project Licence numbers 70/8741 and PP3076489 with approval from the University of Liverpool Animal Welfare and Ethics Review Board. B6 Albino mice (B6N-Tyrc-Brd/BrdCrCrl), a coisogenic mutant of the C57BL/6 strain, were used from a colony maintained at the University of Liverpool colony. Mice were aged 8-11 weeks and male. Mice were housed in individually ventilated GM500 cages with access to food and water ad libitum. A 12-hour light/dark cycle was implemented. Daily checks of animal wellbeing were made and, following surgical intervention, a daily post-operative checklist completed to confirm animal welfare and suitability to remain in the experiment.

### 2.5 Multispectral Optoacoustic Tomography

All MSOT was performed using the inVision 256-TF small animal imaging system (iThera Medical, Germany) [69]. This consists of a tunable (600-1300 nm) optical parametric oscillator (OPO) pumped by an Nd:YAG laser provided 9 ns excitation pulses at a repetition rate of 10 Hz with a wavelength tuning speed of 10 ms. Ten arms of a fibre bundle illuminate a ring of approximately 8 mm in width on the phantom or mouse surface, with fluence below the maximum permissible exposure based on the ANSI safety guidelines. For ultrasound detection, 256 toroidally focused ultrasound transducers with a centre frequency of 5 MHz (60% bandwidth), organized in a concave array of 270-degree angular coverage and a radius of curvature of 4 cm, is used. The phantom holder and the mouse holder are translated using a linear stage to enable imaging of multiple transverse slices.

Images were reconstructed in viewMSOT 4.0.1.34 software (iThera Medical, Germany) with a back-projection algorithm [70] using the BP 4.0 preset to create cross-sectional images. Reconstruction field of view was set to 25 mm (75 μm resolution). Clear, non-absorbing ultrasound gel (Barclay-Swann, UK) was applied to both phantoms and mice for improved ultrasound coupling.

#### 2.5.1 MSOT Phantom Imaging

Phantom inserts were contained within sealed, clear, minimally absorbent straws (Stephensons, UK).

##### 2.5.1.1 Photoacoustic Spectrum Acquisition

Agar phantoms containing two inserts were imaged. One insert contained a solution of the dye in 1X PBS. A second control insert contained PBS alone. The photoacoustic signal of the control straw was subtracted from the photoacoustic signal of the dye straw. The agar phantom was imaged at a water bath temperature of 25°C. Three averaged frames per wavelength were acquired. 45 wavelengths were measured from 680 to 900 nm in steps of 5 nm.

##### 2.5.1.2 Concentration versus Photoacoustic Signal Acquisition

Agar phantoms containing two inserts were imaged. One insert contained a solution of the dye in 1X PBS. A second control insert contained the PBS alone. The photoacoustic signal of the control straw was subtracted from the photoacoustic signal of the dye straw. Six concentrations were measured for each dye (1, 5, 10, 15, 20, 25 μM). The agar phantom was imaged at a water bath temperature of 25°C. Three frames per wavelength imaged were averaged. Measurements were repeated in five positions along the phantom and averaged. 1 wavelength was measured at the peak photoacoustic signal for that dye (700 nm for ABZWCY-HPβCD).

##### 2.5.1.3 Continuous Acquisition

Agar phantoms containing separate inserts of 30 μM ABZWCY-HPβCD or IRDye 800CW in 1X PBS were imaged. The agar phantoms were imaged in a water bath at 34°C. Photoacoustic signal was measured continuously at a single stage position. The MSOT laser was set near the peak photoacoustic signal of the dye (700 nm for ABZWCY-HPβCD and 770 nm for IRDye 800CW). Laser emission was 0.1 Hz. Frames were not averaged.

#### 2.5.2 MSOT Animal Imaging

24 hours prior to imaging, fur was removed with clippers and depilatory cream (Veet Hair Removal Cream 8336076, RB Healthcare, UK) under isoflurane (1-3%) and oxygen (1 L/min) anaesthetic. On imaging day, animals were anaesthetised with isoflurane (1-3%) and oxygen (1 L/min) via a nose cone on a heated mat. The isoflurane dose was titrated to produce a respiratory rate of 1 Hz.

Where the administration of intravenous agents was required, a 30-gauge needle attached to fine bore 0.28 mm polyethylene sterile tubing (Smiths Medical 800/100/100, Fisher Scientific, UK) cut to a length of 33 cm and flushed with 0.9% sterile saline was inserted into the tail vein.

The mouse was placed supine in the MSOT mouse holder with a thin layer of clear non-absorbing ultrasound gel applied (Barclay-Swann, UK). The holder was transferred into the water bath previously heated to 34°C.

#### 2.5.3 MSOT Renal Volume Imaging and Measurement

The kidneys were initially located visually using the MSOT system at 850 nm to identify the renal hila using the live imaging mode. Following 15 minutes acclimatisation to the MSOT water bath, mice were imaged from rostral of the upper renal pole to caudal of the lower renal pole in 1 mm steps in the transverse plane at 850 nm. 10 frames were acquired per stage position and averaged. After image reconstruction, viewMSOT software was used in “Orthogonal View” mode where a region of interest was drawn around the kidney at each slice. From this, the organ volume was calculated by the viewMSOT software.

#### 2.5.4 MSOT Renal Dye Imaging

Following 15 minutes acclimatisation to the MSOT water bath, mice were imaged continuously at a single position for up to 40 minutes with intravenous injection of a dye at 3 minutes. ABZWCY-HPβCD was administered at 30 mg/100 g body weight at a concentration of 150 mg/mL in 0.9% sterile saline and injected over 3 seconds. The volume injected included the administered dose and the volume required to flush the catheter tubing to ensure the full dose was given in a single push without additional flush. The imaging plane was selected visually by identification of the renal hila in MSOT at 850 nm.

Respiratory motion was addressed by titrating the inhaled anaesthetic to achieve a 1 breath per second respiratory rate. MSOT images were then acquired at 10 Hz, and 10 frames per wavelength were averaged to reduce variability due to breathing motion, as previously described [71].

Mice were imaged at the peak absorbance of the relevant contrast agent (700 nm for ABZWCY-HPβCD). Ten frames were measured per wavelength and averaged giving a temporal resolution of 1 second. Time-lapse processing in viewMSOT software was used to quantify the change in dye concentration. A frame 5 seconds before injection was subtracted from the photoacoustic signal as the “zero point”.

On the resulting subtracted frames, regions of interest (ROIs) were drawn for the renal cortex, renal pelvis, aorta, vena cava, and a spinal blood vessel. The ROIs for the renal cortex and medulla were drawn in the dorsal aspect to limit interference from the spleen and liver. The boundary of the spinal blood vessel was identified by a significant peak of contrast immediately following the peak in the aorta.

#### 2.5.5 MSOT Vascular Haemoglobin and Oxygenation Imaging and Measurement

Following 15 minutes acclimatisation to the MSOT water bath, mice were imaged at five wavelengths: 700, 730, 760, 800, and 850 nm. This was measured before any contrast agent was administered. Ten frames per wavelength were measured and averaged. After image reconstruction, images were spectrally unmixed for haemoglobin and oxyhaemoglobin using the linear regression algorithm (viewMSOT, iThera Medical, Germany) [72]. Spectral unmixing algorithms use the total measured spectrum of a pixel and regression to estimate the relative quantities of multiple absorbers in that pixel by their a priori spectra. The same regions of interest representing the aorta, vena cava, and spinal blood vessel used here were copied and pasted from the dye imaging scan.

### 2.6 Statistical Analysis, Normalisation and Mathematical Model Fitting

#### 2.6.1 Statistics

All statistical tests were performed in GraphPad Prism 9.4.0 (GraphPad Software, USA). Paired t test assumes a gaussian distribution and are reported with two-tail P values. Bland-Altman plots were calculated as the difference of A – B vs the average. Pearson correlation coefficients are reported with two-tail P values. Confidence intervals and confidence bands are expressed at the 95% confidence limits.

#### 2.6.2 Normalisation Plots

Plots that were normalised are expressed so that the largest value in each data set is 100% or 1.0 and the value of 0 is 0% or 0.0.

#### 2.6.3 Fitting of Mathematical Models

MATLAB R2020a (MathWorks, USA) was used. Patlak-Rutland (**Equation 3.1**) and Modified Patlak-Rutland (**Equation 3.2**) plots were fit using a 1-degree polynomial with bisquare robust fitting (cftool) and manual exclusion of data points after the initial renal accumulation phase. The integral of the blood concentration against time was calculated by cumulative trapezoidal numerical integration (cumtrapz function). SK-GFR was calculated by the slope of the polynomial fit of the Patlak-Rutland plots multiplied by 0.5 (1 minus the assumed haematocrit [73]) multiplied by the kidney volume measured in MSOT. The sum of the left and right SK-GFR gave the total MSOT GFR.

### 2.7 Transcutaneous Fluorescein Isothiocyanate-sinistrin Measurements and Global GFR Calculation

Mice were anaesthetised by isoflurane (1-3%) and oxygen (1 L/min). A small portion of fur at the dorsum near the loin was shaved and depilated (Veet Hair Removal Cream, RB Healthcare, UK). The adhesive transcutaneous device (MB mini, MediBeacon, Germany) was applied to this region and 3 minutes of baseline readings were recorded. At 3 minutes, 75 μg/g body weight FITC-sinistrin was given by intravenous injection in the tail vein. Following this, mice were recovered from anaesthesia in individual cages. The devices were removed after 90 minutes. FITC-sinistrin curve data was analysed in MB Studio 2 (Medibeacon, Germany) using the 3-compartment fit. A global glomerular filtration rate was calculated from FITC-sinistrin half-life via 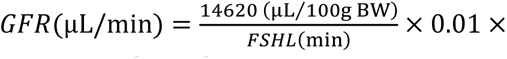 𝐵𝑊(g) where FSHL is FITC-sinistrin half-life and BW is body weight [26,74].

### 2.8 Ischaemia Reperfusion Surgery

Ischaemia reperfusion surgery was performed as previously described [56,75]. Five mice received bilateral injury and five mice received unilateral right-sided injury.

Mice were anaesthetised by isoflurane (1-3%) and oxygen (1 L/min). Mice received analgesia in the form of subcutaneous buprenorphine (1 mg/kg body weight). In addition to this, subcutaneous saline (0.5 mL) and Baytril (70 μL of 10% w/v) were administered. Surgery was performed on a heat pad with feedback from a rectal thermometer to increase and then maintain body temperature at 37°C (Far Infrared Warming Pads, Physiosuite, Kent Scientific, UK). Mice were shaved dorsally (if not already) and prepared with iodine surgical wash. 30 minutes of anaesthetic time elapsed before surgery started to ensure temperature stability of the animal. At the loin, the skin then the muscle was incised using scissors. Abdominal pressure was applied to externalise the kidney via the incision. Forceps were used to expose the kidney blood vessels via blunt dissection. A vascular clamp (Schwartz, Interfocus, Linton) was applied across the vessels at the renal hilum so that a loss of perfusion was visibly identified. A timer was started, and the vascular clamp was removed after either 27.5 or 40 minutes. Reperfusion was confirmed visually. The kidney was gently replaced into the abdominal cavity by wet cotton bud (sterile saline). Muscle then skin was sutured with braided absorbable 6/0 suture (cliniSorb, CliniSut sutures, UK). Mice were then recovered from anaesthesia in a 32°C heat box for 30 minutes.

## 3 Theory

### 3.1 Arterial Input Function

The majority of GFR models require the measurement of the cleared marker’s blood concentration through time which is known as the arterial input function (AIF) [31,34,76]. The AIF can be measured either by repeated blood sampling or during DCE imaging. As mice have a small total blood volume, an imaged AIF is preferred. An imaged AIF can be directly measured by a region of interest (ROI) draw at a blood vessel [76–78].

### 3.2 Models of Glomerular Filtration

#### 3.2.1 Patlak-Rutland Model

The Patlak-Rutland (PR) model is a graphical, two compartment unidirectional kinetic model (Figure 1A, **Equation 3.1**) [35,79–81]. It requires an arterial input function (AIF). It is used in a number of tomography imaging modalities to measure GFR as volume of plasma cleared per time per volume of kidney [31,32,82]. With knowledge of each kidney’s volume, a global GFR and SK-GFR can be calculated. A plot of 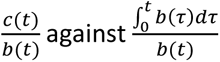 fit with linear regression finds the GFR (the slope) and the fractional vascular volume (FVV, the intercept).

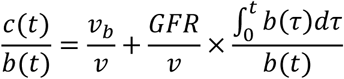

**Figure 1:**
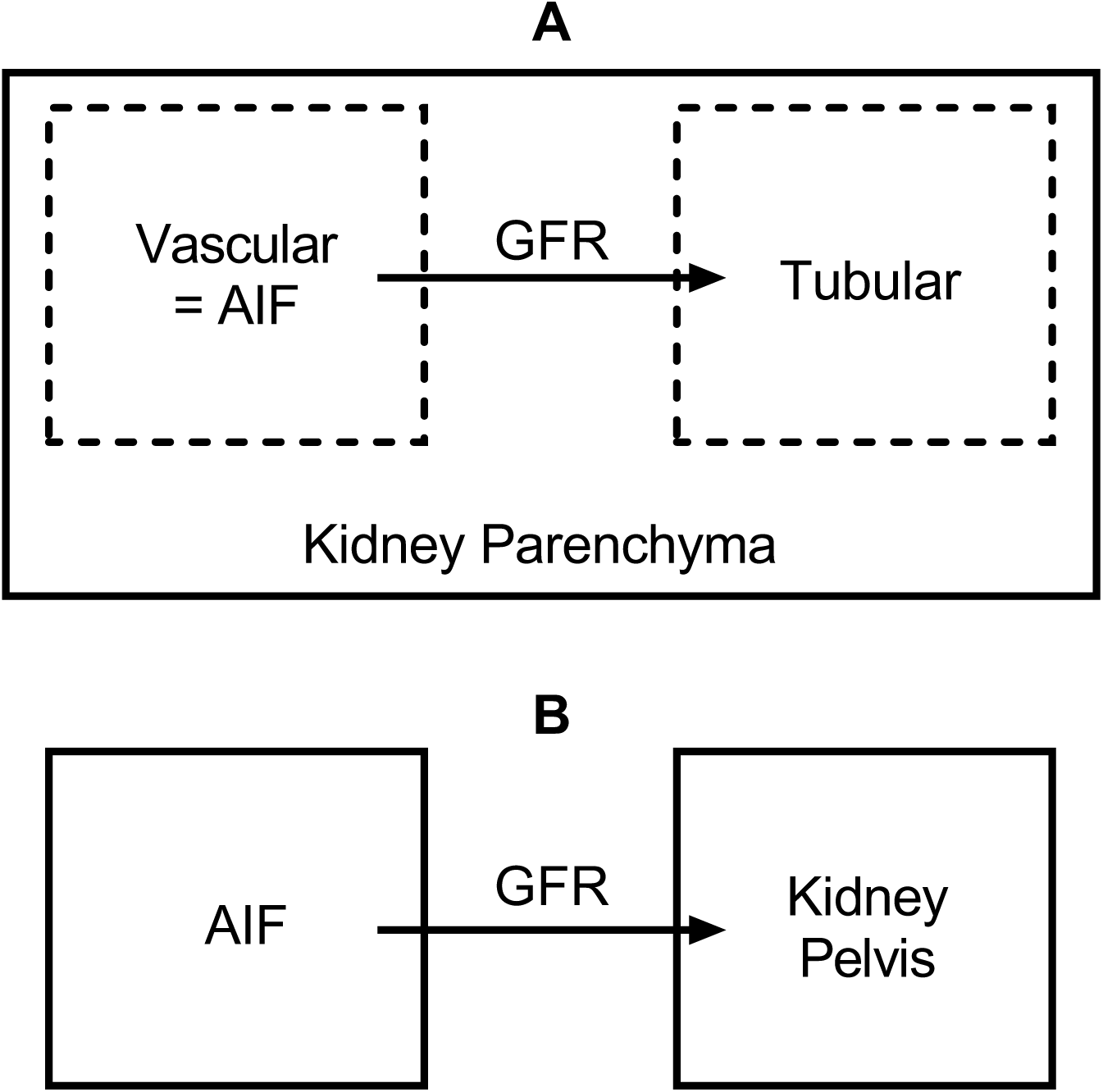
Schematics of pharmacokinetic models of GFR. A, Patlak-Rutland. B, Modified Patlak-Rutland. Solid lines represent a measured region of interest. Dotted lines represent compartments deduced by the model. Arterial Input Function (AIF) is a measured blood vessel e.g., aorta. Glomerular Filtration Rate (GFR).

**Equation 3.1: Patlak-Rutland Model**

t or τ (time), c (kidney cortex concentration), b (blood concentration), v (volume of kidney), v_b_ (blood compartment volume of the kidney)

One of the requirements of the PR model is that the irreversible compartment (the tubular compartment when applied to the kidney) shows negligible outward flow of contrast. This means that the model can only be applied during the uptake phase where dye is still rapidly accumulating in the kidney and tubular outflow is minimal.

#### 3.2.2 Modified Patlak-Rutland Model

Additionally, we have developed a modified version of the PR model (MPR, Figure 1B, Equation 3.2). Due to the rapid renal cortex outflow seen with ABZWCY-HPβCD, application of a standard PR model is limited by a short uptake phase. The MPR model compensates for this by using the renal pelvis where the uptake phase lasts for several minutes. As the renal pelvis is avascular, the term that represents the fractional vascular volume is absent.

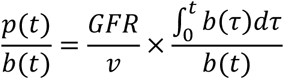

**Equation 3.2: Modified Patlak-Rutland Model**

t (time), p (kidney pelvis concentration), b (blood concentration), GFR (glomerular filtration rate), v (volume of the kidney)

## 4 Results

### 4.1 Phantom Experiment Results

ABZWCY-HPβCD was examined in tissue mimicking agar phantoms in MSOT to characterise the photoacoustic spectrum and the relationship between dye concentration and photoacoustic signal intensity. The photoacoustic spectrum and optical absorbance spectrum of ABZWCY-HPβCD are similar (Figure 2A). We found a slight left-shift in the peak (695 nm from 705 nm) and a broader absorbance curve from the photoacoustic spectrum. A linear relationship between ABZWCY-HPβCD concentration and photoacoustic signal was identified when imaged at 700nm in MSOT (Figure 2B, R^2^ 0.9611).

**Figure 2:**
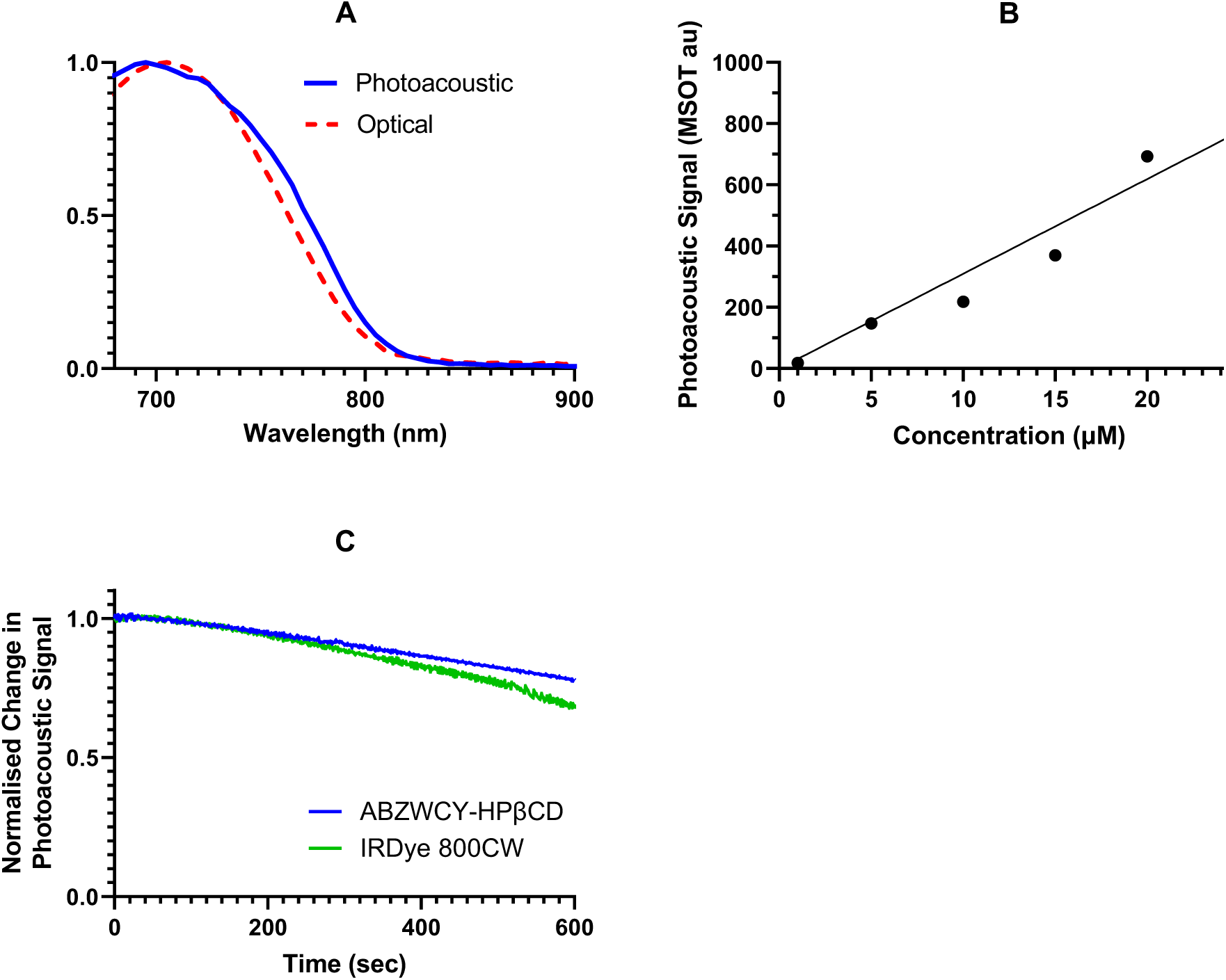
ABZWCY-HPβCD. A, Normalised spectra of ABZWCY-HPβCD: photoacoustic spectrum in MSOT phantom and optical absorbance in microplate reader. B, Photoacoustic signal as a function of ABZWCY-HPβCD concentration in MSOT phantom. C, Normalised change in photoacoustic signal of ABZWCY-HPβCD and IRDye 800CW with continuous MSOT imaging in phantoms.

As NIR dyes are known to undergo photodegradation [83–85], we examined whether there was evidence of a change in photoacoustic intensity with continual NIR laser irradiation in MSOT. This is important because a change in photoacoustic intensity should reflect changes in concentration that are due to clearance rather than degradation. ABZWCY-HPβCD and IRDye 800CW were compared in phantom concerning their photoacoustic signal stability on continuous imaging at their peak absorbing wavelength. ABZWCY-HPβCD showed progressive loss of photoacoustic signal when imaged continually in MSOT in agar phantom (Figure 2C). However, this compared favorably with IRDye 800CW.

### 4.2 ABZWCY-HPβCD DCE MSOT in Healthy Mice

Seven mice were examined. FITC-sinistrin clearance was quantified using a transcutaneous device to calculate a global GFR. Following this, mice were imaged with MSOT to measure ABZWCY-HPβCD clearance.

#### 4.2.1 ABZWCY-HPβCD Distribution

The dye rapidly peaks in the aorta, vena cava, and renal vasculature (Figure 3B). After this, the dye accumulates in the renal cortex (Figure 3C) and then in the renal pelvis (Figure 3D). Dye also accumulates in and then clears from the skin and soft tissues (Appendix A: Supplementary Figure 1).

**Figure 3:**
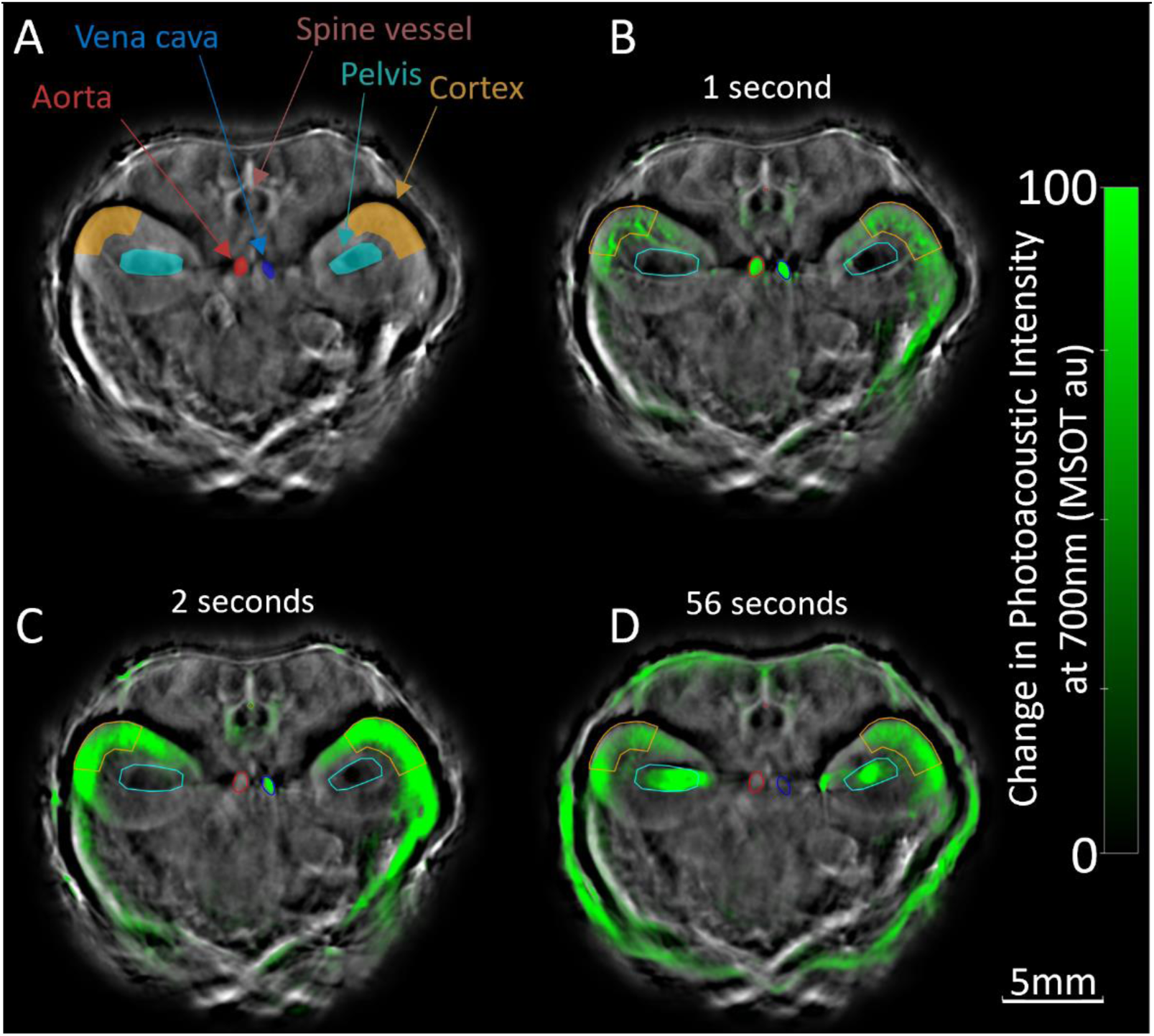
Typical ABZWCY-HPβCD distribution after IV administration measured by DCE MSOT. A, Regions of interest measured for models. B, Vascular peak. C, Renal cortex Peak. D, Renal pelvis peak.

#### 4.2.2 Arterial Input Function

A region of interest was drawn at the aorta and the photoacoustic signal intensity over time was measured. The pre-injection signal was subtracted to calculate the change in signal. In all cases, a signal below pre-injection levels was seen immediately after an initial peak in the aorta (Figure 4A). Over the course of the imaging session, the signal slowly drifts back to baseline. This “negative” signal significantly impacts our ability to use the aorta as an AIF because it is not possible to have a negative concentration.

**Figure 4:**
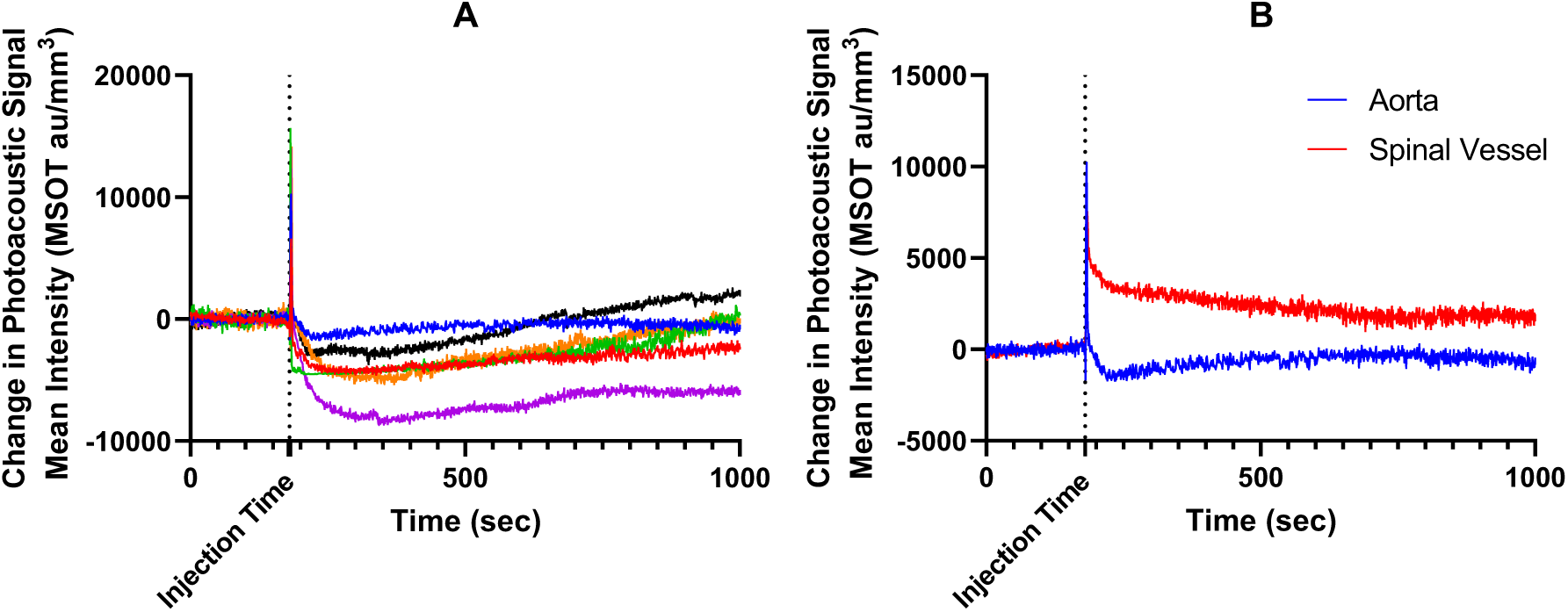
Selection of a Suitable Vessel for AIF in MSOT with ABZWCY-HPβCD. A, Examples of the change in mouse aorta photoacoustic signal following injection of ABZWCY-HPβCD where each coloured line represents a different mouse. B, Example comparing aorta and spinal blood vessel photoacoustic signal following injection of ABZWCY-HPβCD.

To assess a more superficial blood vessel that would make a suitable AIF in DCE MSOT, we measured a spinal blood vessel. A ∼0.2 mm diameter circular region of interest was drawn in the spine where a hyperintense signal was seen in the frames immediately following ABZWCY-HPβCD injection. The change in photoacoustic signal intensity over time was measured in the same way as the aorta. When this was compared with the aorta, the change in photoacoustic signal over time in the spinal blood vessel did not show any dip below pre-injection signal in any cases (Figure 4B). These curves showed a peak immediately after the aorta followed by an apparent multi-phase exponential decay.

#### 4.2.3 Modelling GFR

The PR model doesn’t account for the flow of dye from the tubules to the renal pelvis and bladder (Figure 1A). As such, a requirement of the model is to fit only the uptake of the contrast in the kidney. ABZWCY-HPβCD has shown a significantly short uptake phase in the renal cortex (Figure 5A). The initial uptake in the renal cortex occurs after 3 seconds. While there is a smaller increase that could possibly be part of the uptake phase after 30 seconds, this is dramatically earlier than Jiang et al saw with their renally cleared gold nanocluster, Au25(SG)18 [50]. The renal pelvis shows a longer uptake phase with ABZWCY-HPβCD (Figure 5A). Because of this, the renal pelvis was used as part of our Modified Patlak-Rutland model (Figure 1B). Examples of the time activity curves and Patlak-Rutland plots for these two models can be seen in Figure 5B-E.

**Figure 5:**
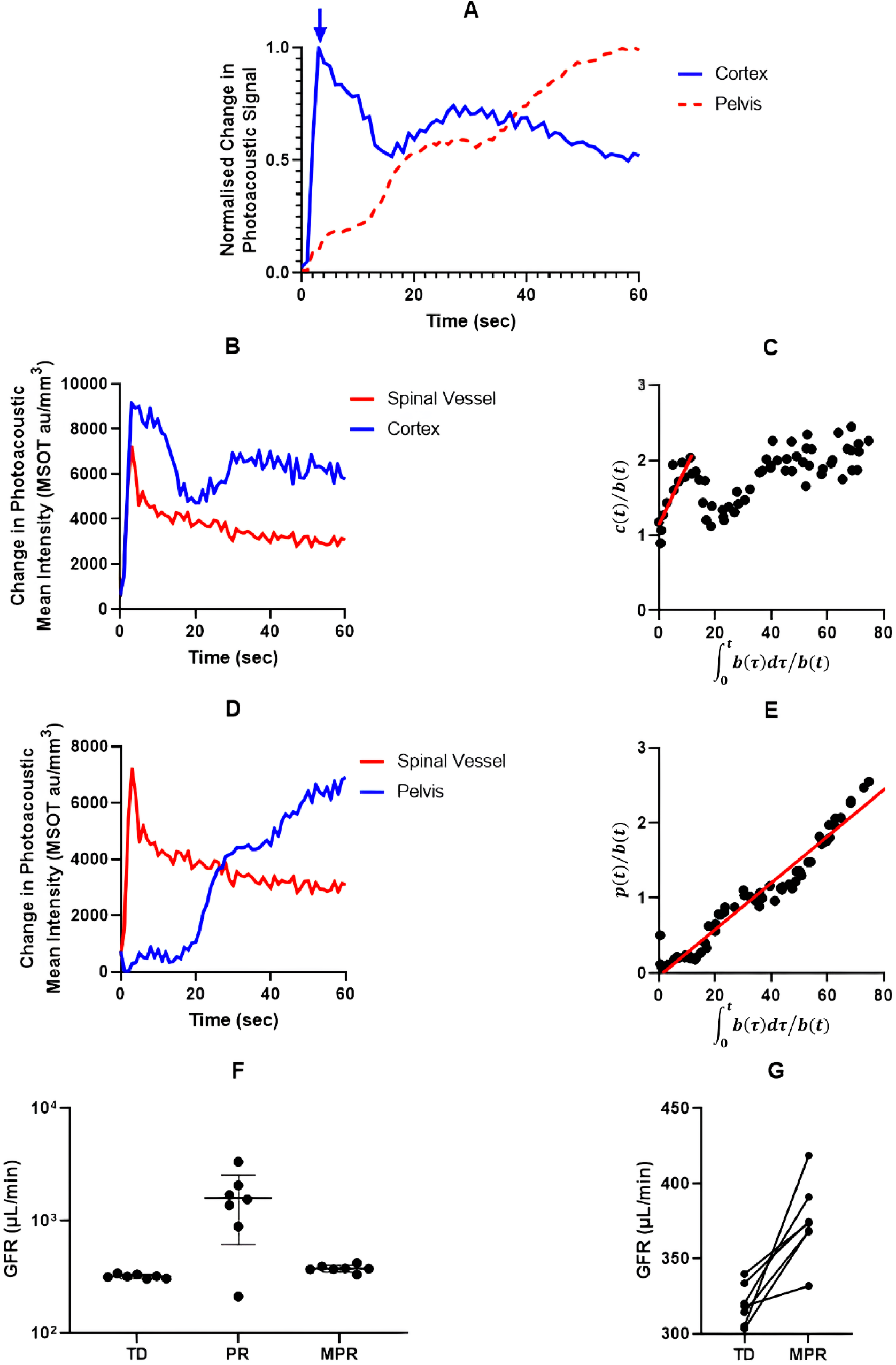
Time activity curves (TACs) of ABZWCY-HPβCD, example Patlak-Rutland plots and GFRs in healthy mice. A, TACs in MSOT following ABZWCY-HPβCD injection (arrow marks end of uptake phase in the cortex). B, TACs for ROIs applied to PR model with spinal blood vessel AIF. C, PR plot for B. D, TACs for ROIs applied to MPR model with spinal blood vessel AIF. E, MPR plot for D. F, GFR values obtained with PR and MPR models of DCE MSOT GFR compared with the Transcutaneous Device (TD) (clearance of FITC-sinistrin). G, Correlation plot between TD GFR and DCE MSOT GFR MPR model, indicating a strong correlation (Pearson r = 0.9195, P < 0.0001).

Using the kidney volumes measured in MSOT and the slope of the Patlak-Rutland plots, global GFRs were calculated from the DCE MSOT data using a spinal vessel AIF. In the healthy animals, a standard Patlak-Rutland model overestimated the GFR significantly in all but one case (Figure 5F) while the Modified Patlak-Rutland model gave a far more reasonable approximation of the global GFR measured by determining the clearance of FITC-sinistrin with a transcutaneous device (Figure 5F&G).

### 4.3 ABZWCY-HPβCD DCE MSOT following IRI

A further ten mice underwent examination. A 27.5-minute surgical IRI was performed: five mice received bilateral IRI while five received unilateral IRI. Mice had their transcutaneous device GFR and MSOT ABZWCY-HPβCD clearance measured on three occasions: prior to injury, post-op day 1, and post-op day 3. These were compared with a global GFR calculated by transcutaneous device-measured FITC-sinistrin clearance.

To further characterise the spinal blood vessel AIF, photoacoustic signal was measured at multiple wavelengths (700, 730, 760, 800, 850 nm) prior to ABZWCY-HPβCD injection at baseline imaging. Then spectral unmixing algorithms were applied to quantify haemoglobin, oxyhaemoglobin, and haemoglobin saturation. The concentrations of these absorbers within the spine blood vessel region of interest were compared with the aorta and vena cava (Figure 6). The quantification of these absorbers were similar between the spinal blood vessel and the other two vessels measured suggesting a vascular structure had been correctly identified.

**Figure 6:**
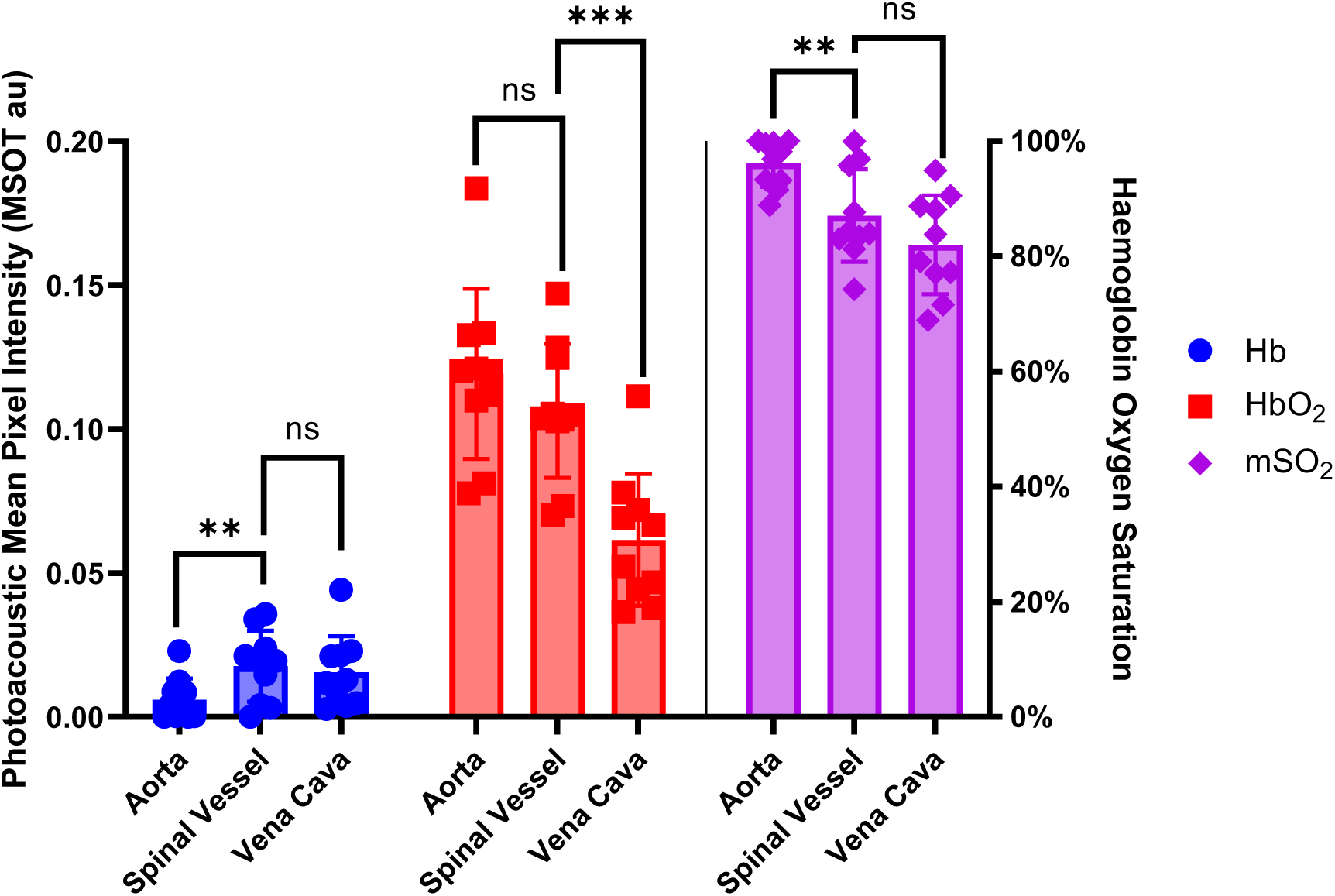
MSOT spectral unmixing of for deoxyhaemoglobin, oxyhaemoglobin, and haemoglobin oxygen saturation in the spinal blood vessel, aorta, and vena cava of mice. Hb (Deoxyhaemoglobin), HbO2 (Oxyhaemoglobin), mSO2 (Haemoglobin Oxygen Saturation). ns P >0.05, * P ≤0.05, ** P ≤0.01, *** P ≤0.001, **** P ≤0.0001. Bars are the standard deviation.

Our Modified Patlak-Rutland (MPR) model was applied and examined for correlation with transcutaneous device GFR at all 30 time points (Figure 7). Global GFR calculated by MPR ABZWCY-HPβCD DCE MSOT showed good correlation with FITC-sinistrin transcutaneous device GFR across a range of GFRs with a Pearson r of 0.9195 (P = <0.0001, Figure 7A). 95% confidence intervals were narrow (0.8362 to 0.9613). Bland-Altman plot showed that the Modified Patlak-Rutland method overestimates GFR in healthy mice while underestimating GFR following IRI (Figure 7B). The 95% limits of agreement were -117.7 to 124.3.

**Figure 7:**
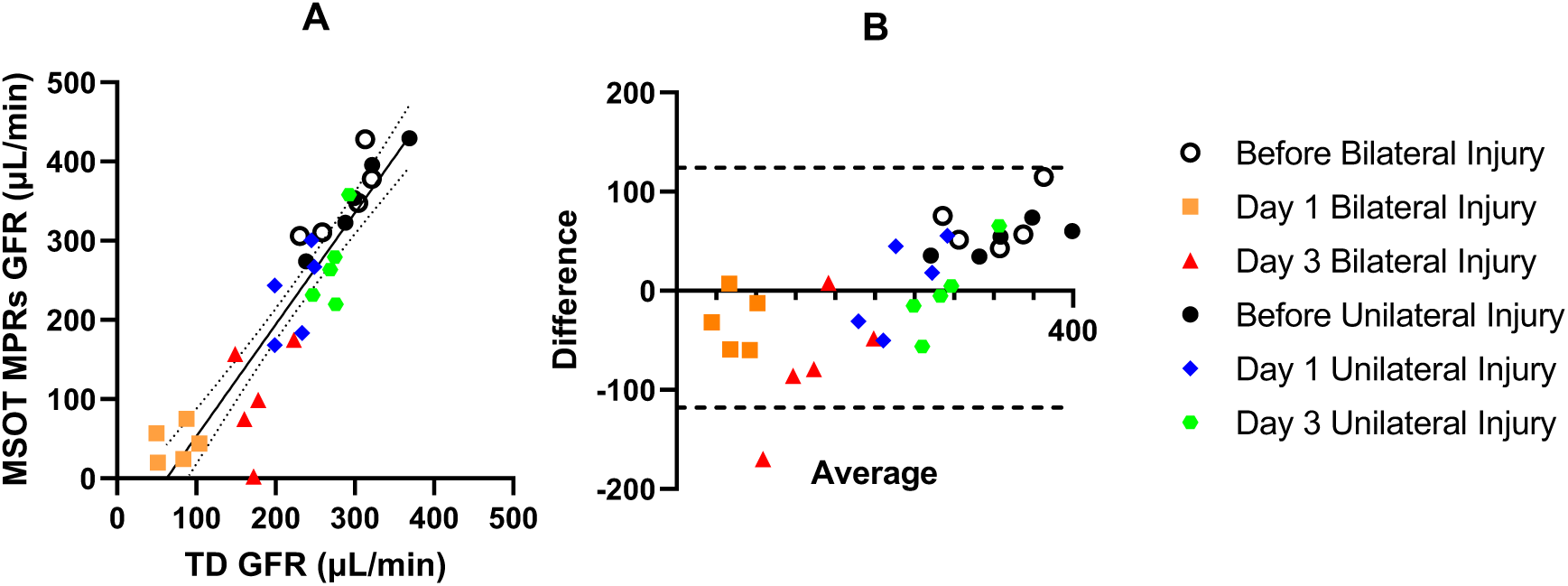
Correlation between MSOT MPR GFR and Transcutaneous Device GFR. A, Transcutaneous Device versus MPR GFR, Pearson r = 0.9195 (0.8362 to 0.9613, R^2^ 0.8454). B, Bland-Altman plot of A. Transcutaneous Device, MPR (Modified Patlak-Rutland with spinal blood vessel AIF). Solid line is a simple linear regression fit. Dotted lines are the 95% confidence bands. Dashed lines are 95% Limits of Agreement.

The SK-GFR calculated using the MPR method was used to detect differences in renal function between two kidneys in a model of unilateral IRI (Figure 8). SK-GFR was calculated using the MPR model for mice receiving both the bilateral and unilateral IRI.

**Figure 8:**
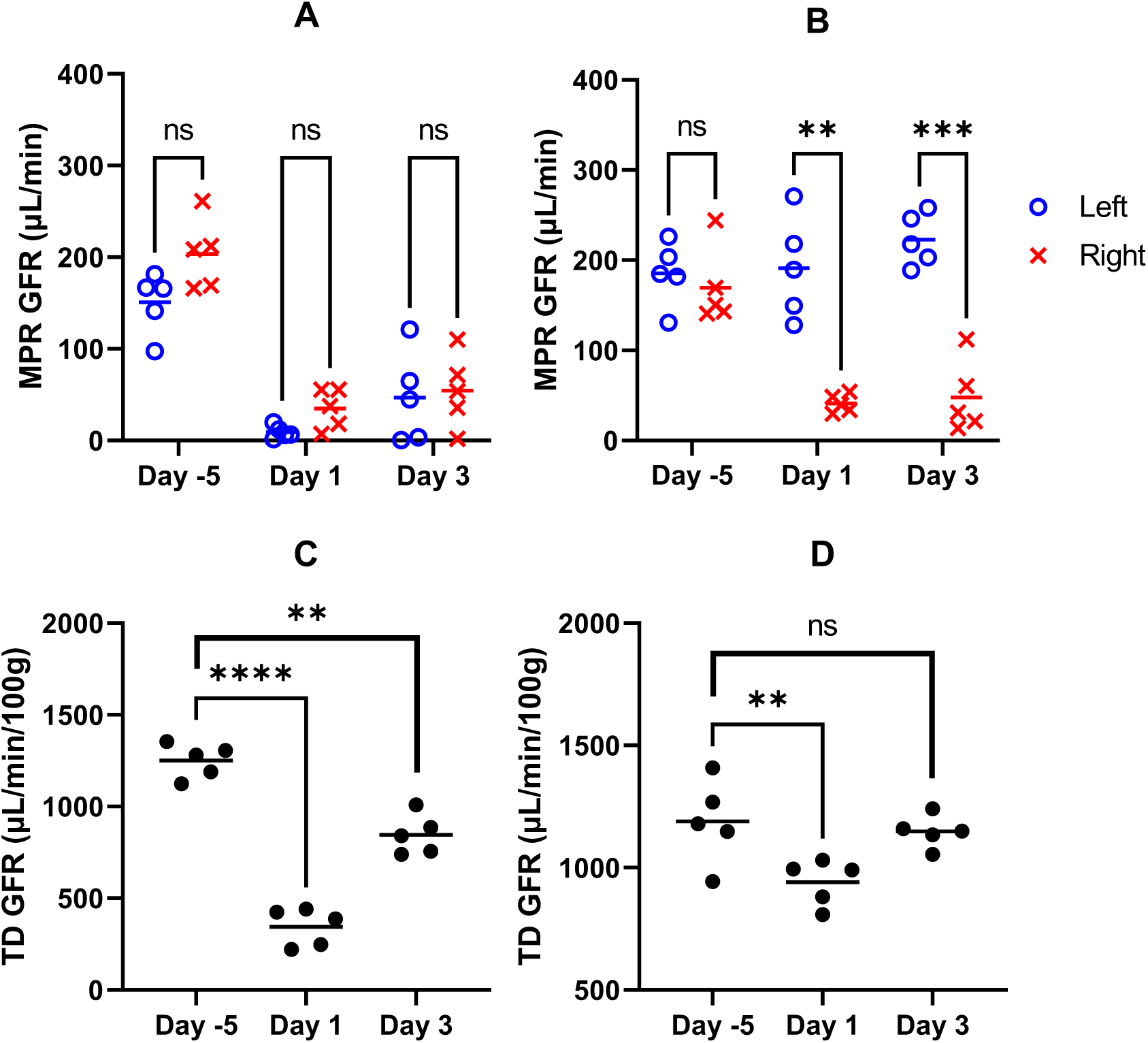
MSOT and Transcutaneous Device GFR in Ischaemia Reperfusion Injury. A, MSOT Modified Patlak SK-GFR in Bilateral injury. B, MSOT Modified Patlak SK-GFR in Unilateral injury. C, Transcutaneous Device GFR in bilateral injury. D, Transcutaneous Device GFR in unilateral injury. Horizontal line is the mean. ns P >0.05, * P ≤0.05, ** P ≤0.01, *** P ≤0.001, **** P ≤0.0001.

At baseline, the right kidney SK-GFR was moderately higher than the left SK-GFR in the bilateral injury group (Figure 8A). However, this was not significant. On day 1 post injury the GFR of both kidneys dropped similarly with the right kidney function remaining slightly higher than left in the bilateral injury group. Day 3 saw some recovery. There was no statistical difference between left and right SK-GFR at any point in the bilateral injury group.

In the unilateral injury group, the left kidney SK-GFR was mildly higher than the right kidney prior to injury, though this was not significant (Figure 8B). On day 1 and 3 post injury the right kidney was significantly impaired while the left kidney function remained similar. There was a statistical difference between left and right after injury on both injury days (Day 1 P = <0.005, Day 3 P = <0.001). Figures 8C and 8D show the GFR obtained with the transcutaneous device (FITC-sinistrin clearance) in the bilateral and unilateral IRI model, respectively. In the unilateral model, the GFR at day 3 is not significantly different from baseline GFR. This is expected given that the transcutaneous method measures global GFR.

## 5 Discussion and Conclusion

Here we report on DCE MSOT using ABZWCY-HPβCD to measure GFR and SK-GFR in mice before and after surgical IRI. We found a good correlation between DCE MSOT GFR and GFR measured by determining FITC-sinistrin clearance with a transcutaneous device when a MPR model and a spinal blood vessel AIF was used. This method successfully delineates unilateral kidney injury from the internal control when measuring SK-GFR.

Pharmacokinetic modelling of GFR in DCE imaging requires an AIF. While blood sampling is considered the gold standard AIF measurement [34], mice have small blood volumes meaning repeated sampling can lead to hypovolaemia resulting in poor animal welfare and impaired GFR results. As such, an imaged AIF is preferred. Following ABZWCY-HPβCD intravenous injection in MSOT, the aorta shows an initial peak in photoacoustic signal followed by a fall below pre-injection levels. This “negative” signal makes the use of the aorta as an AIF impractical because it would represent a negative concentration of dye which is a physical impossibility. The intensity of photoacoustic signal is the result of the amount of delivered light, the efficiency with which the absorber converts heats to a pressure related expansion, the concentration of the absorber, and the acoustic coupling [67]. The amount of light delivered to a point is dependent on the point’s depth as well as the absorbance and scattering of the light in the tissues superficial to this [86]. As the aorta is a deep structure near the centre of the mouse, it is susceptible to a reduction in light delivered particularly as absorbance increases in the periphery of the mouse due to dye accumulation there. This hypothesis is supported by the progressive return to baseline over the course of the imaging session as dye clears from the periphery. Algorithms that can account for the delivery of light to different points in a sample are under active development [67,87,88]. Jiang et al did not report any issue with the aorta as the AIF using their renally cleared gold nanoclusters to measure SK-GFR in DCE MSOT via a PR model [50]. This may be due to the different natures of the contrast agents. ABZWCY-HPβCD may show greater peripheral accumulation increasing its chances of impairing light delivery to the deep tissues. ABZWCY-HPβCD also shows lower photoacoustic signal generation than their gold nanoclusters at similar concentrations and so is less detectable at lower concentrations.

Using a superficial blood vessel in the spine as the AIF in ABZWCY-HPβCD DCE MSOT showed more typical time activity curves. As this spinal blood vessel is at a similar depth to the kidneys, a similar amount of light could be expected to be delivered to these regions. Spectral unmixing of these regions showed haemaglobin, oxyhaemaglobin, and haemaglobin saturations similar to the aorta and vena cava confirming that a region containing blood had been selected. It is unclear which vessel this represents. Due to the location, it may be the posteromedial spinal vein. While this vessel would not contain arterial blood, it did not appear to negatively impact our results. This may be due to the animal’s rapid heart rate resulting in similar arterial and venous concentrations.

The outflow of ABZWCY-HPβCD from the renal cortex is rapid. We observed photoacoustic signal in the cortex drop six to ten seconds after injection. While a second less intense peak is frequently seen in the 40 seconds following the first peak, the general trajectory of the signal is downward. This means that the uptake phase in the cortex is relatively short. A short uptake phase in the renal cortex impairs the application of a standard PR model. This model does not account for the outflow of marker from the irreversible compartment. The initial increase in the cortex is likely more representative of the dye in the vascular compartment of the kidney. The dye that has been filtered into the tubules has no opportunity to accumulate in the cortex ROI because the outflow rate is so high. This leads to the overestimation of GFR by a standard Patlak-Rutland model. Small molecule NIR dyes are known to be quickly excreted and frequently are cleared by the kidneys [51,55]. As such, a rapid outflow of ABZWCY-HPβCD is in keeping with this. The fast clearance of small molecule NIR dyes may contribute to their lower risk of toxicity. We observed a far longer uptake phase in the renal pelvis, and this allowed the implementation of our MPR model and likely explains the improved correlation with sinistrin-measured GFR. The MPR model overestimates the GFR in healthy animals and underestimates the GFR following IRI. As a comparison, DCE MRI has shown both overestimation and underestimation of GFR depending on the model and acquisition used [31,42–45,89]. Following IRI surgery, identification of the renal pelvis in MSOT is more challenging due to oedema, sutures, and changes in positioning of the kidneys [56]. We have also noted a reduction in the native contrast between the cortex and the medulla following IRI. Also, the rate at which ABZWCY-HPβCD moves through the tubules is possibly reduced following IRI. As the pelvis is not the site of filtration, the MPR model relies on the rapid outflow of dye from the cortex to the pelvis to make the tubule to pelvis transit time negligible and the calculated rate representative of GFR. If the flow through the tubules is reduced in injury, the tubule to pelvis transit time may no longer be insignificant. Furthermore, the amount of ABZWCY-HPβCD accumulating in the renal pelvis would be significantly lower in injury and such a low concentration in the renal pelvis may be less detectable by MSOT leading to an underestimation of GFR.

The MPR model was able to detect a significant difference between the injured and uninjured kidney in a unilateral 27.5-minute IRI. Transcutaneous device global GFR only detected a significant difference on day 1 post unilateral injury. The MPR model showed an increase in the mean SK-GFR of the uninjured kidney on day 3. This compensatory increase in the control kidney may explain why the Transcutaneous Device global GFR has normalised on day 3. Contralateral kidney compensation is seen in models of unilateral nephrectomy in rodents where there is an increase in renal blood flow in the remaining kidney [90–92]. Changes in renal blood flow occur within 15 minutes and an increase in SK-GFR occurs as soon as 48-hours following nephrectomy [91,92]. Acute SK-GFR compensation following nephrectomy appears to be driven by a reduced production of renin, reduced stimulation of angiotensin II type 1 receptors, and increased renal blood flow produced by decreased intra-renal vascular resistance [91]. However, IRI is associated with an upregulation of angiotensin II type 1 receptors with a maintained production of renin [93]. This achieves a higher SK-GFR by increasing intraglomerular pressure. Angiotensin II type 1 antagonism has been shown to reduce progression to CKD after IRI induced AKI independent of systemic blood pressure control [94]. We did not measure renin–angiotensin–aldosterone system (RAAS) products nor blood pressure, but this would be a valuable area of investigation. There may be an association between MSOT MPR-detected contralateral filtration compensation, changes in RAAS products, systemic blood pressure, and intra-renal vascular resistance.

ABZWCY-HPβCD has not previously been measured by MSOT and we have demonstrated the ability to detect this dye both in phantoms and mice. It has been shown that DCE MSOT using ABZWCY-HPβCD and a MPR model using a spinal blood vessel as the arterial input function correlates well with GFR calculated using FITC-sinistrin and a transcutaneous device. This provides preclinical researchers with the opportunity to measure SK-GFR at a simultaneously high spatial and temporal resolution without ionising radiation. This technique can facilitate a greater understanding of injury and recovery in injury models via an internal control kidney. ABZWCY-HPβCD is able to measure SK-GFR in MSOT due to the dye’s high urinary recovery, low protein binding, and absence of tubular secretion [28] improving over the current NIR dye standard, IRDye 800CW. The zwitterionic NIR dye, ZW800-1, which has been found to be a promising fluorophore for assessing renal function in mice using NIR fluorimetry [95], might also prove to be effective for assessing renal function using MSOT. However, it was beyond the scope of this study to directly compare ZW800-1 with ABZWCY-HPβCD.

## Author Contribution Statement

James Littlewood drafted the manuscript and performed MSOT imaging, analysis, and modelling. Rachel Harwood performed surgical procedures. James Littlewood, Rachel Harwood, and Katherine Triviño-Cepeda performed transcutaneous GFR measurements. Srishti Vajpayee synthesised ABZWCY-HPβCD. Rossana Perciaccante, Norbert Gretz, Jack Sharkey, Thomas Sardella, Neal Burton, Tim Delving, Arthur Taylor, Bettina Wilm, Rachel Bearon, and Patricia Murray provided supervision and expert advice.

## Supporting information

Dye clearance movie

## Acknowledgements

We acknowledge use of the Centre for Preclinical Imaging (CPI) imaging facilities provided by Liverpool Shared Research Facilities, Faculty of Health and Life Sciences, University of Liverpool. We acknowledge support for animal maintenance and breeding of the strain by the Biomedical Sciences Unit at the University of Liverpool. This project has received funding from the European Union’s Horizon 2020 research and innovation programme under the Marie Sklodowska-Curie grant agreement No. 813839.

## Data Availability

All data is freely available at Zenodo via the following DOIs: 10.5281/zenodo.10500280

10.5281/zenodo.10500297

10.5281/zenodo.10500308

10.5281/zenodo.10650178

10.5281/zenodo.10650192

## Vitae

**James Littlewood** is a specialist registrar in renal medicine. He received his BMBS from Peninsula Medical School in 2012. He received his PhD in physiology from the University of Liverpool in 2022. His research focused on the use of optoacoustic tomography to assess renal function. He was sponsored by iThera Medical and the European Union’s Horizon 2020 scheme.

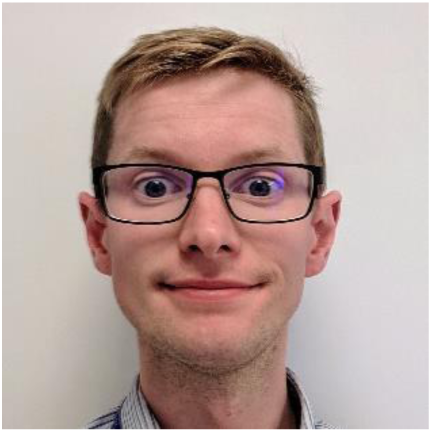

**Rachel Harwood** is an Academic Clinical Lecturer at the University of Liverpool and a Paediatric Surgery Registrar at Alder Hey Children’s Hospital. She undertook a PhD looking at regenerative therapies for acute kidney disease and used novel imaging techniques to define the degree of kidney injury. Rachel’s ongoing research interests include the translation of regenerative medicine therapies into clinical practice including their safety profile and efficacy.

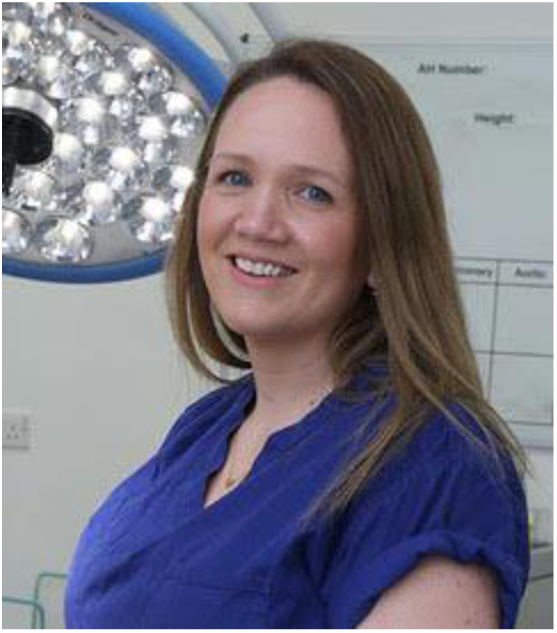

**Thomas Sardella** is currently an application specialist at iThera Medical. He got his PhD degree from Glasgow University on spinal cord injury research and carried out his postdoctoral work at Glasgow University on pain pathways in the spinal cord. His research interests are focused on clinical and preclinical applications in photoacoustics.

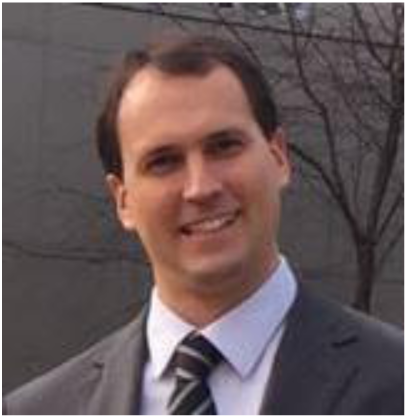

**Neal Burton** has worked at iThera for 13 years. As the head of the Applications department, he supervises the global team of Application Specialists. The Applications team at iThera is primarily responsible for training and supporting existing customers in experimental design and data analysis, identifying new applications, and performing demonstrations of PAI for prospective customers. NCB received his undergraduate degree from Washington University in St. Louis in 2001, and his PhD in Environmental Health Sciences, from the Johns Hopkins Bloomberg School of Public Health, Division of Toxicology, in 2008.

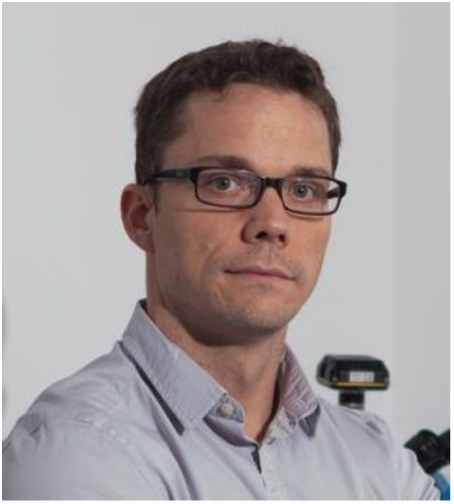

**Norbert Gretz** was the head of the Medical Research Center of the Medical Faculty Mannheim, University of Heidelberg, and the managing director of the Institute for Medical Technology, a combined institute of the Medical Faculty Mannheim, University of Heidelberg, and the University of Applied Sciences Mannheim. He became senior physician at the University Hospital Mannheim. He was appointed professor for experimental medicine at the University of Heidelberg. His key research areas were photobiomodulation, chronic wounds, kidney diseases, stem-cell therapy, optical-tissue clearing, gene-expression profiling, and big data.

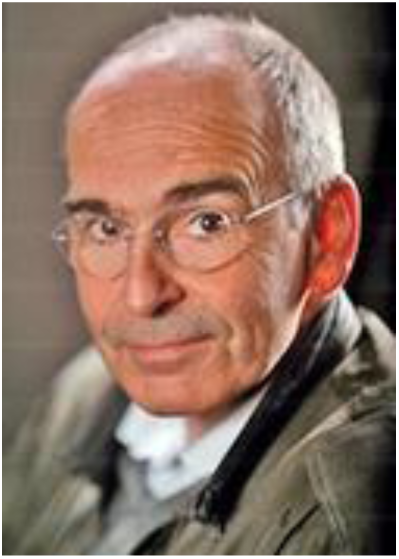

**Rachel Bearon** is Executive Dean of the Faculty of Natural, Mathematical & Engineering Sciences at King’s College London. She was previously Head of the Department of Mathematical Sciences at the University of Liverpool, where she was also appointed Professor in Mathematical Biology. She leads research that applies mathematics to health challenges, focusing on the spatial and temporal dynamics of biological systems, ranging from bacterial chemotaxis, cancer cell motility and phytoplankton in turbulence, to modelling cell-signalling pathways, intracellular protein dynamics and drug transport.

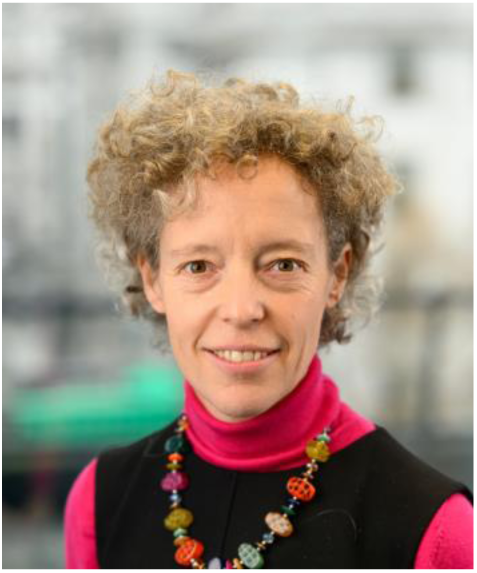

**Rossana Perciaccante** is Research & Development Manager at Cyanagen srl. After earning her MSc in Chemistry from Bologna University (2000), she worked as Junior Researcher at GSK (UK). After her Ph.D. degree in Organic Chemistry in 2006 from Bologna University, she joined Cyanagen where she worked as researcher of the National Centre of Excellence LATEMAR focusing on the development of fluorescent sensors for genomic, post-genomic and biomedicine. She is co-author of several scientific papers and patents in the field of chemistry and biological sciences. She participates in national and international scientific research programs. Her scientific interests include the development of new fluorescent and chemiluminescent probes for imaging and molecular diagnostics.

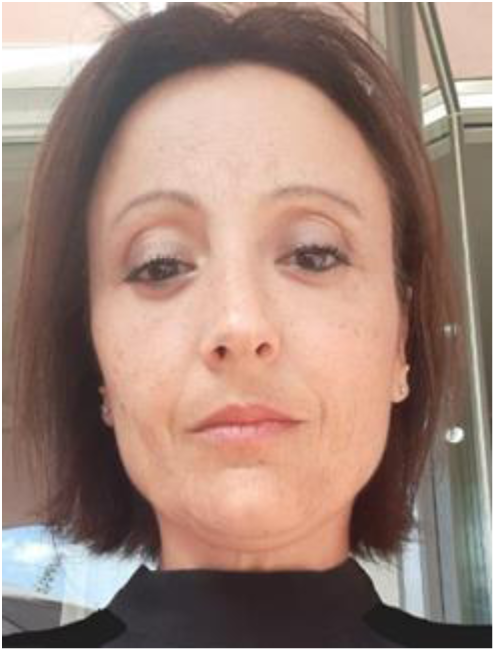

**Jack Sharkey** is a subject matter expert as part of the in vivo imaging team at Revvity. He completed his PhD at the University of Liverpool where he investigated micro RNAs as potential mechanistic and translational biomarkers of kidney injury. He subsequently carried out a postdoc in cell based regenerative medicine therapies and continues to have an interest in preclinical imaging in the field of regenerative medicine.

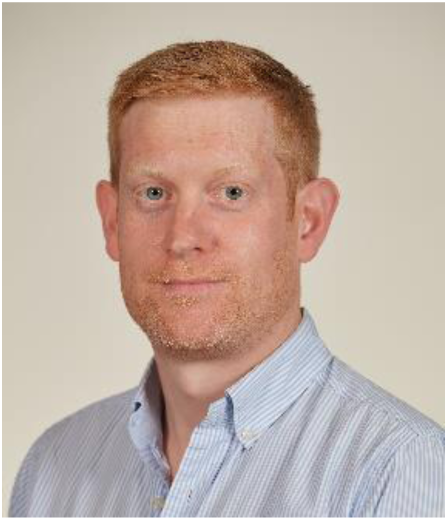

**Arthur Taylor** is a Research Associate at the University of Liverpool, UK. His interests lie in the development of novel methods for molecular imaging. Before moving to Liverpool, Arthur obtained his PhD from the Dresden University of Technology (Germany) and a double MSc degree from the Delft University of Technology (The Netherlands) and the Dresden University of Technology (Germany). He completed his bachelor’s degree at the Federal University of Santa Catarina (Brazil).

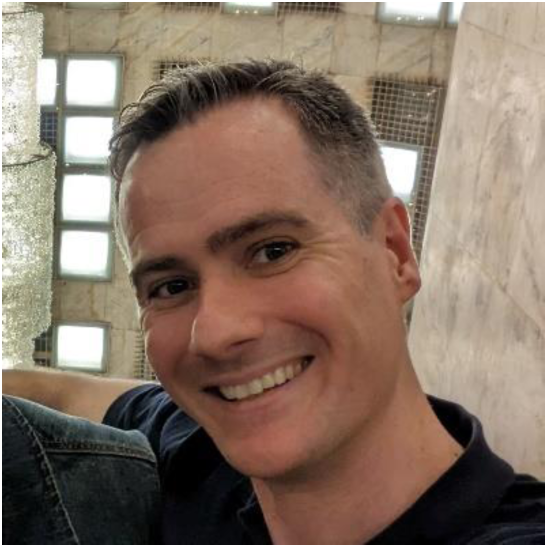

**Katherine Trivino-Cepeda** is currently pursuing a Ph.D. degree in Molecular Physiology and Cell Signaling at the University of Liverpool in England. She holds a Bachelor’s degree in Biotechnology Engineering from the University of the Armed Forces (ESPE) in Ecuador and she has completed an Erasmus Mundus Joint Master’s program in Medical Science from Uppsala University in Sweden and Medical and Pharmaceutical Drug Innovation from the University of Groningen in the Netherlands. Katherine is interested in translating research into the clinical practice and accelerating the development of innovative therapeutic solutions for unmet medical needs.

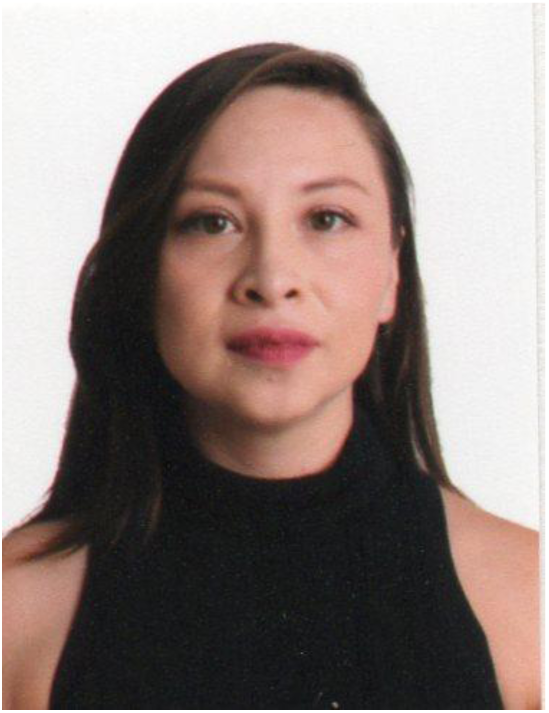

**Srishti Vajpayee** obtained her B.Sc degree in Biomedical Sciences from University of Delhi (New Delhi, India) in 2017 and her M.Sc. degree in the same field from Radboud University (Nijmegen, The Netherlands) in 2019 specialising in molecular life sciences. In August 2019 she received European Union’s H2020 Maria Skłodowska-Curie fellowship to carry out her industry-academia Ph.D. at Cyanagen Srl in Italy and Medical Faculty Mannheim, University of Heidelberg in Germany. She completed her Ph.D. degree in 2022 under the supervision of Dr. Rossana Perciaccante and Prof. Norbert Gretz, focusing on the development of novel fluorescent agents for a non-invasive assessment of kidney function and kidney imaging.

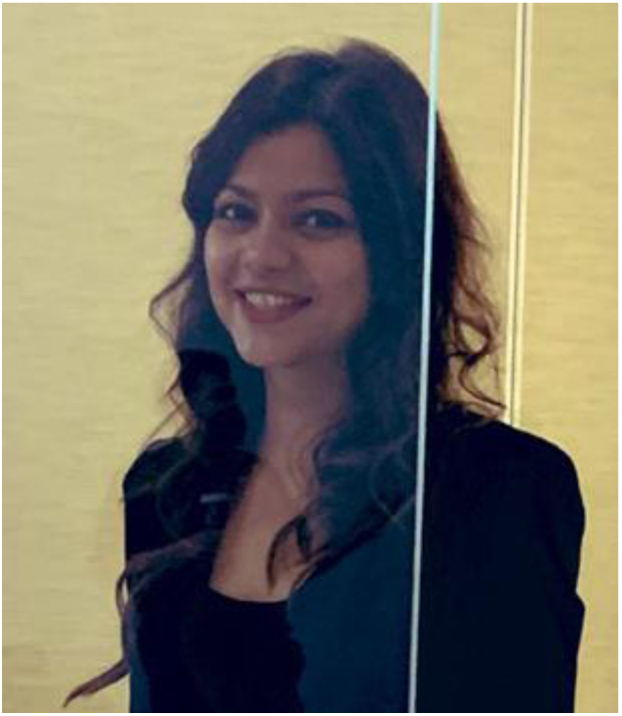

**Bettina Wilm** is a Senior Lecturer at the University of Liverpool. Over the last 10 years, she has successfully led research projects using rodent kidney injury models to assess cells as regenerative medicine therapies. As part of this aim, her team has developed new methods of detecting the kidney damage physiologically and via novel preclinical imaging techniques. The development of these technologies will be crucial in establishing the efficacy of regenerative medicine therapies in rodent models, on the route to clinical applications.

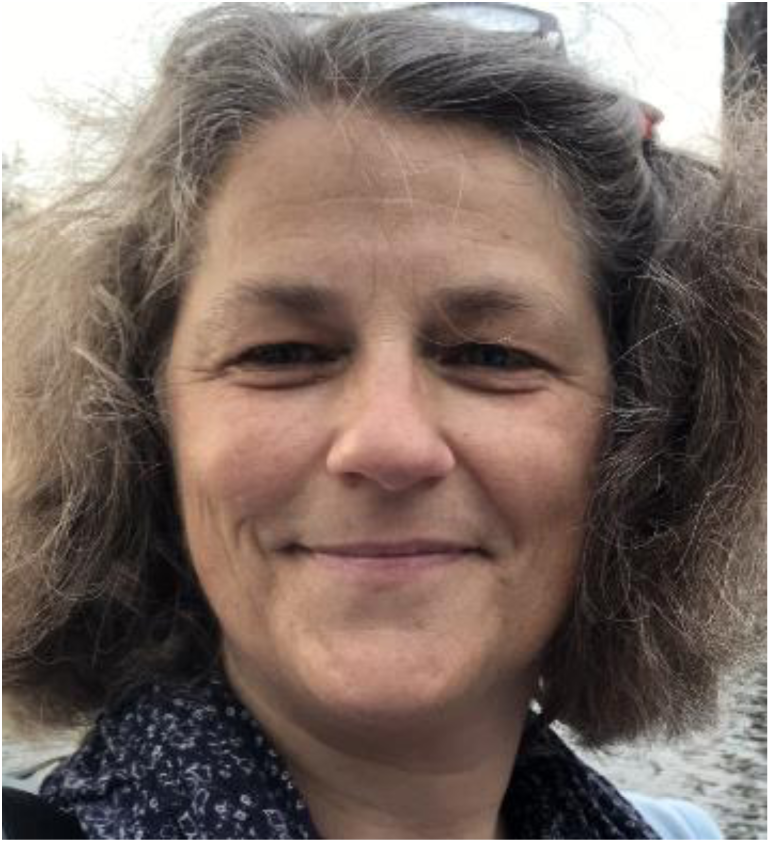

**Patricia Murray** is a professor of stem cell biology and regenerative medicine at the University of Liverpool. She has a long-standing interest in investigating the therapeutic potential of renal regenerative medicine. This includes developing non-invasive imaging techniques to determine the fate of administered cell therapies, and new imaging approaches to monitor kidney function over time in small animals.

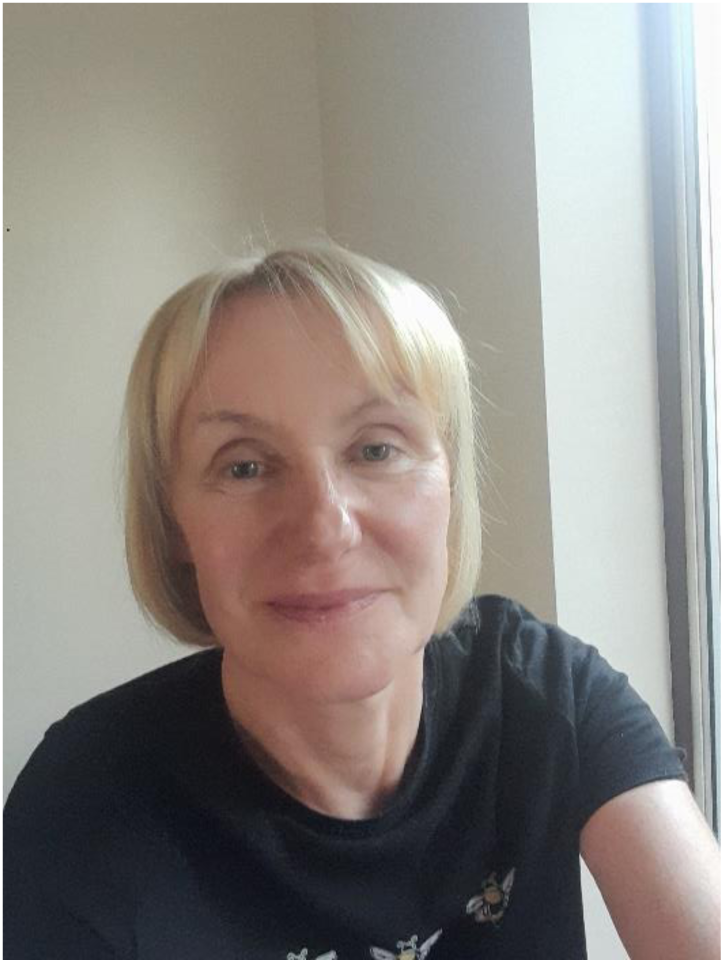

