## Supplementary material for "Modelling Glomerular Filtration using Multispectral Optoacoustic Tomography and a Novel Near-infrared Dye": Dye clearance movie

### Slide 1
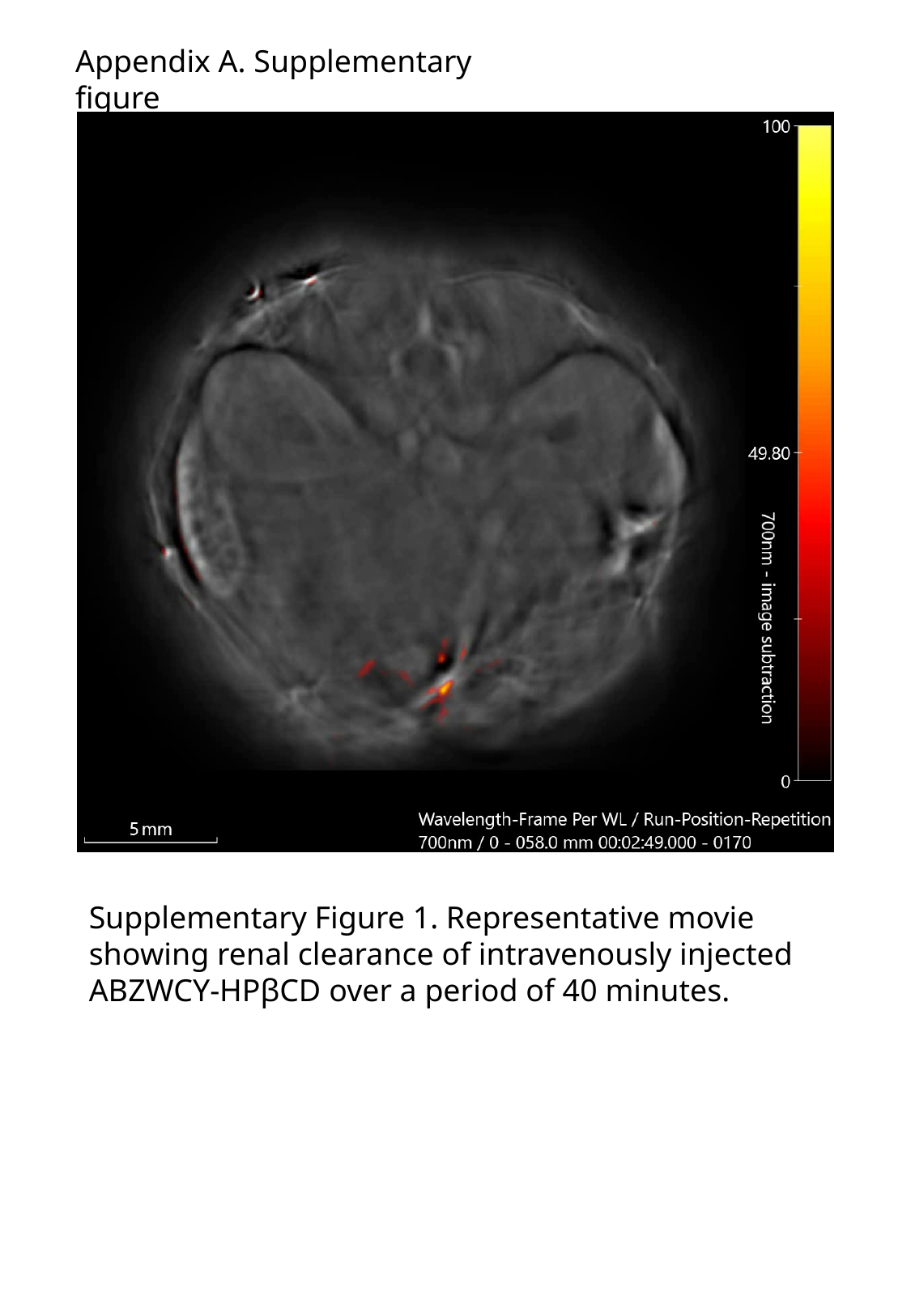

Appendix A. Supplementary figure
Supplementary Figure 1. Representative movie showing renal clearance of intravenously injected ABZWCY-HPβCD over a period of 40 minutes.
